# STRipy: a graphical application for enhanced genotyping of pathogenic short tandem repeats in sequencing data

**DOI:** 10.1101/2021.06.13.448220

**Authors:** Andreas Halman, Egor Dolzhenko, Alicia Oshlack

**Affiliations:** Peter MacCallum Cancer Centre, Melbourne, Victoria 3000, Australia; Sir Peter MacCallum Department of Oncology, The University of Melbourne, Victoria 3010, Australia; Murdoch Children’s Research Institute, Royal Children’s Hospital, Parkville, Victoria 3052, Australia; Florey Department of Neuroscience and Mental Health, The University of Melbourne, Parkville, Victoria 3010, Australia; School of Natural Sciences and Health, Tallinn University, 10120 Tallinn, Estonia; Illumina Inc., 5200 Illumina Way, San Diego, CA, USA; School of BioSciences, University of Melbourne, Parkville, Victoria 3052, Australia

## Abstract

Short tandem repeats (STRs) are highly polymorphic with high mutation rates and expansions of STRs have been implicated as the causal variant in diseases. The application of genome sequencing in patients has recently allowed many new discoveries with over 50 disease causing loci known to date. There are several tools which allow genotyping of STRs from high-throughput sequencing (HTS) data. However, running these tools out of the box only allow around half of the known disease-causing loci to be genotyped, with lengths often limited to either read or fragment length which is less than the pathogenic cut-off for some diseases. While analysis tools can be customised to genotype extra loci, this requires proficiency in bioinformatics to set up, use, and analyse the resulting data, limiting their widespread usage by other researchers and clinicians.

To address these issues, we have created a new software called STRipy that has an intuitive graphical interface and requires no specific skills for usage, thus significantly simplifying detection of STRs expansions from human HTS data. STRipy is able to target all known disease-causing STRs with genotyping performed with an established tool, ExpansionHunter, that is incorporated into the software. We have created additional functionality into STRipy to work with long alleles exceeding the fragment length.

STRipy was validated using over 60 thousand simulated samples and was shown to work on whole genome sequencing of biological samples with pathogenic variants. Finally, we have used STRipy to acquire genotypes of pathogenic loci for thousands of samples from various populations which are provided to the user along with the data from the literature to assist with results interpretation. We believe the simplicity and breadth of STRipy will increase the testing of STR diseases in current datasets resulting in further diagnoses of rare diseases caused by STRs expansions.

## Introduction

Short tandem repeats (STRs) are sequences consisting of repeated, short motifs up to 6 bp long. They comprise about 3% of the human genome [1]. Over 50 human diseases are reported to be caused by expansions in the number of repeats at specific loci in the human genome and around one fifth of these have been discovered in the past five years [2]. Many of the recently discovered disease-causing STRs are different to the vast majority of previous findings as the pathogenic motif was found to be absent in the reference genome which is based on healthy individuals. These include repeats that cause different types of familial adult myoclonic epilepsies (FAMEs) [3–6], spinocerebellar ataxia 37 (SCA37) [7] and cerebellar ataxia neuropathy vestibular areflexia syndrome (CANVAS) [8].

Analysis of STRs from short-read sequencing data is not straightforward due to the low complexity and non-uniqueness of sequences from these regions. Therefore STR analysis has often been ignored in medical sequencing studies [9]. However, advances in sequencing technology and repeat-aware software over the past decade has resulted in several bioinformatics tools built for genotyping STRs, some of which can also estimate repeat length for alleles longer than fragment length, such as ExpansionHunter [10] and GangSTR [11]. ExpansionHunter is a tool created specifically for targeted genotyping analysis and has been used successfully on a large-scale analysis of rare diseases, detecting alleles which are shown to be well correlated with PCR-based assays [12]. The latest version (4.0.2) of the tool uses a variant catalogue containing information which allows the estimation of STR genotypes in 28 loci that are known to cause a disease. However, only two of these diseases can report genotypes that are longer than the sequencing fragment length. In addition, the provided variant catalogue does not contain information for detecting any of the recently discovered loci where a pathogenic motif was inserted between or next to the endogenous STRs.

STRs genotyping tools are generally command-line programs which fit well into bioinformatic pipelines, but are not intuitive to set up and use on their own without relevant skills, limiting their everyday application to the analysis of sequencing data. Here, we introduce a new free and open-source software with graphical interface called STRipy, which is an easy to use program built to detect all known pathogenic STRs to date. The first aspect of STRipy is the curated database of all discovered disease causing STRs with supporting genomic and pathogenic information. STRipy then incorporates ExpansionHunter [10] into its framework as the backend genotyper and has increased the number of defined loci from 28 to 55, to include all currently known disease-causing loci. We additionally integrated a tool named REViewer [13] to our software to visualise haplotype sequences and reads aligned on them, which also enables the visualisation of any interruptions in the STR locus. STRipy has added functionality to enable genotyping of alleles that are longer than the sequencing read fragment length and also reports the number of pure pathogenic motifs in reads.

Moreover, STRipy genotyping was applied to two and a half thousand individuals from the 1000 genome cohort in order to provide the distribution of population level genotypes for each locus. This, together with information from literature, is provided to improve assessment of pathogenicity of genotype in a sample. The resulting online database of disease-causing STRs with population data for all loci provides a valuable resource for all scientists in the field. Ultimately, STRipy aims to improve the clinical diagnoses of rare diseases and patients’ care by increasing detection of STRs expansions which can now be performed easily by all researchers and clinicians.

## Design and implementation

### STRipy’s Client

STRipy is implemented in Python 3 and consists of two parts – Client and Server. The Client is the main part of the software which a user interacts with through a graphical interface. It allows the genotyping of one locus at a time to provide quick results and avoid incidental findings. The whole genotyping process is streamlined, only requiring the user to specify from a drop-down menu: a locus or disease associated with the locus to target; a BAM or CRAM file of aligned and indexed paired-end short-read sequencing data; and the genome assembly the BAM/CRAM file was aligned to. The software can also autodetect the genome which further simplifies the process for a user. STRipy’s autodetection algorithm scans the sequences before and after the targeted STR locus in the analysed sample and determines the most likely genome. If no matches are found, the user would need to select it manually. The type of analysis can be chosen to be either ‘Quick’ or ‘Extended’, with the latter suitable for genotyping alleles longer than the fragment length. Finally, a user can choose whether to run the analysis in their own computer/internal network (Local) or remote server (Cloud). Upon clicking on the button to genotype the sample, a cascade of processes will be executed.

STRipy uses a catalogue containing variant information such as the reference coordinate and sequence of the repeat unit required for the bioinformatics analysis. At first, STRipy’s Client extracts out reads from in and around the targeted STR locus. In the case of the ‘Extended’ analysis, STRipy determines the location of mis-aligned mates of the reads that are aligned next to the STR locus and extracts out those mates as well as all reads which are fully made of the pathogenic repeat unit, if they exist (Fig 1). These locations are specified as off-target regions for ExpansionHunter to extend the genotyping estimation above the fragment length. Additionally, STRipy determines the number of pathogenic motifs in reads which can be useful when the allele contains interruptions or is made of repeats different from the endogenous ones. The presence or lack of the pathogenic repeat unit is critical information to the user when analysing replaced and nested type of repeats as ExpansionHunter will also count similar (endogenous) repeat units into the length of the reported genotype even when there is no pathogenic motif present. Finally, a BAM file with an anonymised file name is created, containing the extracted reads, which is then indexed and forwarded to STRipy’s Server for genotyping.

**Fig 1.**
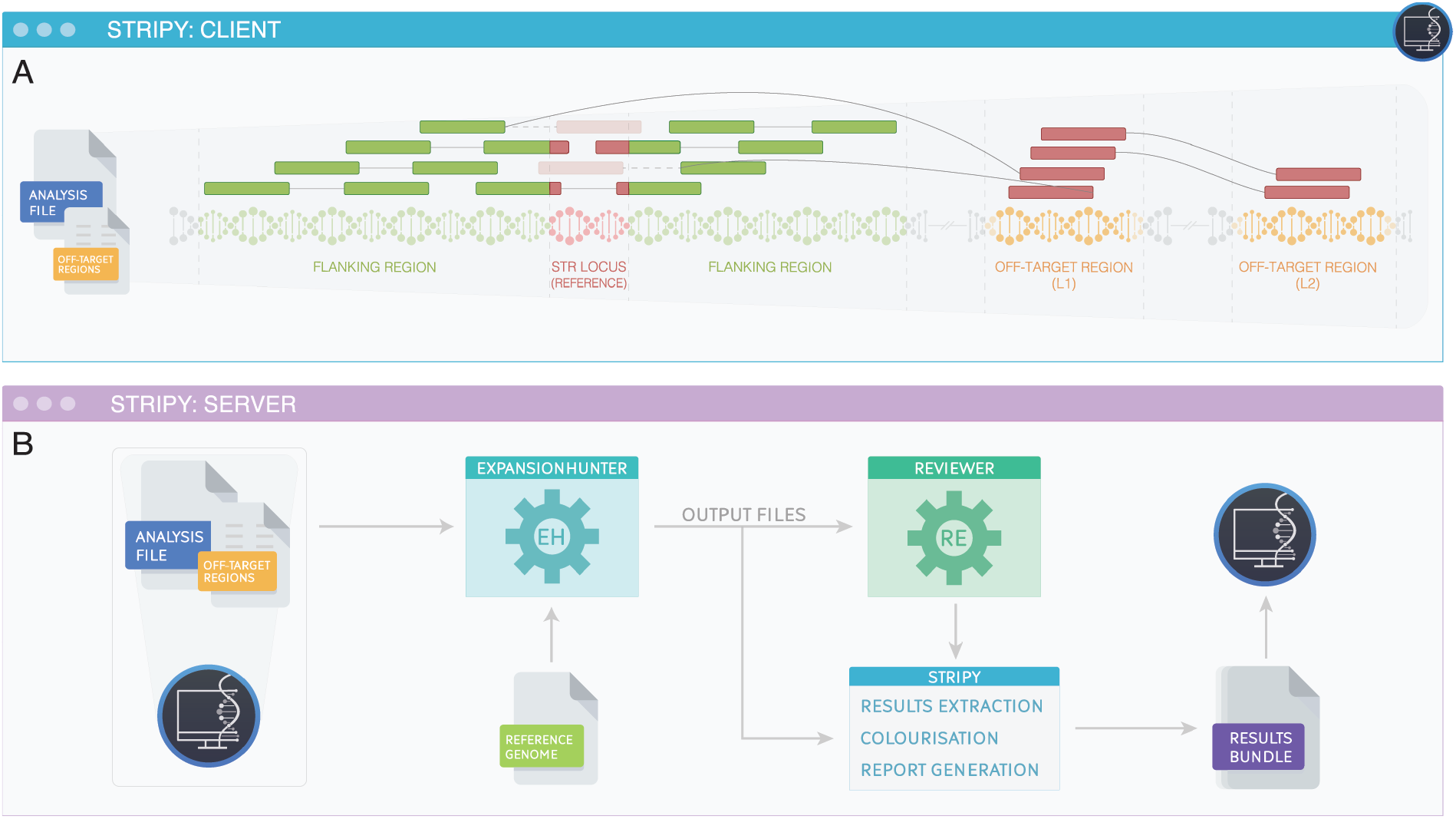
Overview of STRipy’s method. (A) From the sequencing file, reads within or overlapping a flanking region (2 kb by default) each side of the STR locus (marked in green) will be extracted out in STRipy’s Client on local computer. By using the ‘Quick’ analysis, flanking reads containing the STR region (marked in both green and red) and fully repeated reads (marked in red) that are aligned elsewhere but where one read in a pair is in next or overlapping STR locus will be included into the analysis file. When using the ‘Extended’ analysis, besides extracting out mates of the reads aligned next to the STR locus, STRipy uses their locations and tries to find more fully repeated and misaligned reads in level 1 (L1) off-target regions. If they exist, then these reads will be extracted out as well with their mates, some of which can be aligned to another, level 2 (L2) off-target region. The resulting analysis ready file for ‘Extended’ analysis includes reads from the region surrounding the reference STR locus and reads found from L1 and L2 off-target regions, which is then forwarded to the STRipy’s listening server (B) where it will be genotyped by the ExpansionHunter using STRipy’s determined off-target regions. In case of the ‘Quick’ analysis, no off-target regions are specified. Read visualisations are created with REViewer, colourised and then combined with genotyping results obtained from ExpansionHunter and returned to the STRipy’s client along with the generated PDF report.

### STRipy’s Server

STRipy’s Server listens for inputs from the Client and performs several tasks after receiving the data. First, the received BAM file will be genotyped by ExpansionHunter [10] using variant information (reference coordinate and repeat unit) which was acquired from literature and manually verified by us. In case of the ‘Extended’ analysis, off-target regions found by STRipy’s Client will be forwarded to ExpansionHunter to enable genotyping of long alleles. This approach is different from using ExpansionHunter outside of STRipy as we are using the data to determine off-target regions for each sample individually, in contrast to using pre-defined regions for all samples. After genotyping, ExpansionHunter’s output files will be analysed by the tool REViewer [13] which creates read alignment visualisations for the STR region. Read visualisations is an important part of the results section for two reasons. Firstly, it allows visual assessment of how well the genotype is supported by reads aligned to the locus. Secondly, this shows the most likely sequence for both alleles, including the presence of interruptions within repeats which can be clinically important, for example by impacting the timing of disease onset [14]. In the end, genotyping results from the ExpansionHunter along with the visualisations of read alignments are returned back to the STRipy’s Client after going through colourisation process (Fig 2). The Client then presents the genotyping results accompanied by repeat specific information obtained from literature as well as the populationwide data we have generated to improve assessment of the results.

**Fig 2.**
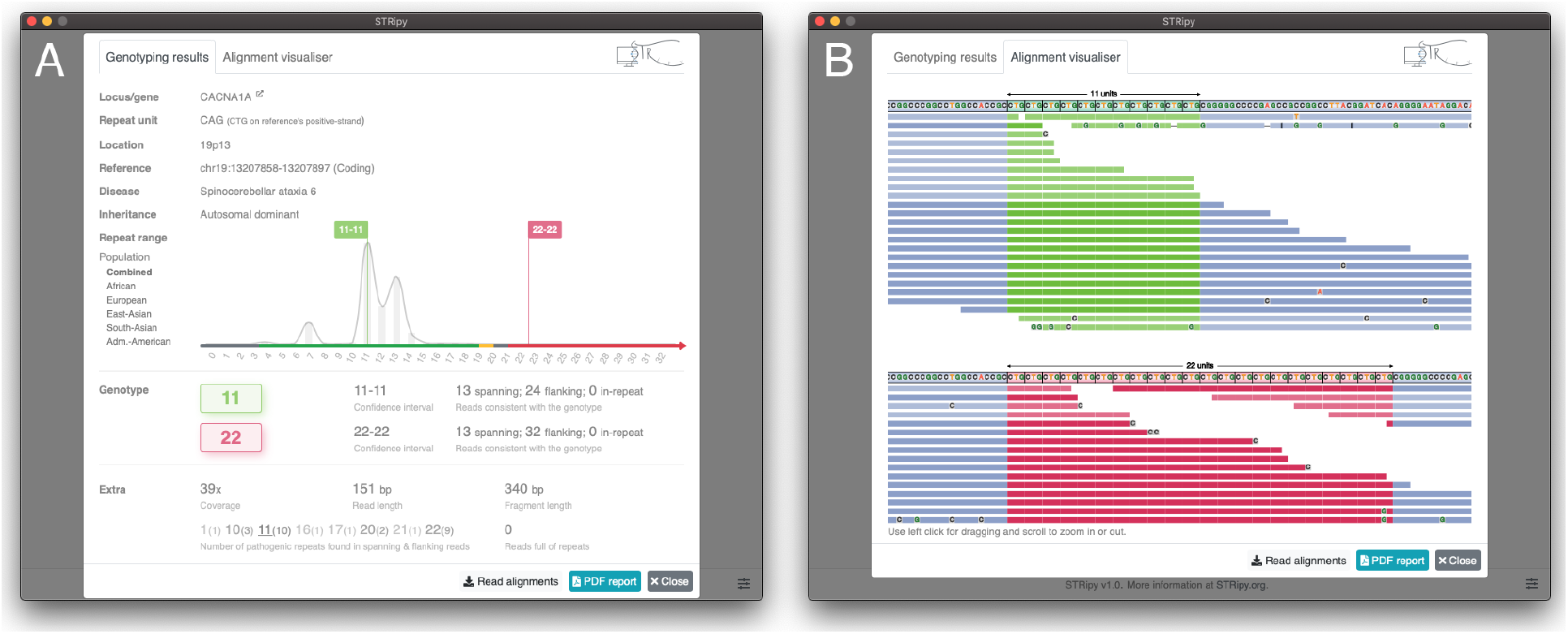
Screen captures of STRipy’s Client. (A) the upper half of the screen contains information obtained from the literature for the locus (genomic location, associated disease, inheritance and pathogenic repeat ranges displayed as a colourised X-axis line). The reference position and repeat unit is also from literature, but reviewed and modified by us to be in 0-based coordinate system and uses motif corresponding to this region. The population-wide data distribution for the locus is shown on top of the X-axis line, which can be changed to represent data for each of the super-populations separately (African, European, East or South Asian and Admixed-Americans). On the bottom half of the screen, the genotyping results for the sample from ExpansionHunter are displayed under the “Genotype” section as well as coverage, read and fragment length. Below that, are two fields containing results from STRipy’s algorithm that report the presence of the pathogenic repeat unit, such as in the flanking reads and total number of reads full of repeats. Read alignments can be found under the Alignment visualizer tab (B). A PDF report can be saved by clicking the corresponding button in the bottom.

STRipy’s Server can be easily installed on Linux and macOS operating system, requiring Python 3 with svglib, regex, pyramid and reportlab library. The server needs to have access to the reference genome(s) as well as to Samtools [15], ExpansionHunter and REViewer executables. To further simplify the process of genotyping STRs, we provide a free STRipy’s Server (Cloud) from our end where all the tools and main reference genomes are provided. The Cloud has been configured for maximum privacy and it does not store sequencing information nor any other information of the sample (including genotyping results) or any user identifying information. By using the Cloud, a user does not have to set up a server on their own, which makes genotyping STRs an exceptionally easy process as there is no installation or set-up when using the compiled STRipy’s Client that contains Python 3 as well as all the required libraries (pysam, pywebview, requests, numpy, regex). STRipy’s Client works both on macOS and Linux operating systems.

### STRipy’s STRs database

We created the most comprehensive and up-to-date STRs database for use by STRipy and to make available for more general purposes. This database contains variant information for each locus assisting with bioinformatics analysis and results interpretation. We performed a comprehensive search in literature to determine all disease-causing STRs loci reported so far, collected the variant information, manually confirmed and adjusted all reference coordinates as well as repeat unit’s sequence for different genome versions (S1 Table 1–3; Supplementary information). In total we determined 55 loci in 52 genes where repeat expansions are reported to be causative for 58 diseases. This will be updated new disease causing STR expansions as they are discovered and reported.

We classified our list of loci into three groups based on the repeat type: standard (34 loci), imperfect GCNs (12 loci) and replaced/nested (9 loci). Standard repeats are considered to be the ones where one specific repeat unit (for example, CAG or CCG) is in healthy individuals and its expansion becomes pathogenic at larger numbers of repeats (S1 Table 1). Imperfect GCN type refers to repeats which encodes for the amino acid alanine, but the sequence can be composed of different, but synonymous, repeat units – either GCA, GCG, GCC or GCT (S1 Table 2). Lastly, replaced/nested type of repeats were determined to be ones where the pathogenic motif is not present in the reference genome and usually not found in healthy individuals (S1 Table 3). Out of the nine diseases discovered so far, only one (CANVAS) is caused by replaced type repeats, where a stretch made of repeat units with one sequence (AAAAG) is replaced with a sequence composed of another motif (AAGGG). All the other diseases in this group are caused by nested type of repeats where a stretch made of the pathogenic motif (such as TTTCA) is inserted between or next to the non-pathogenic stretch of endogenous repeats (such as TTTTA).

Finally, we used STRipy to genotype all 55 loci in 2,504 samples from the DRAGEN reanalysis of 1000 Genomes project dataset [16] to obtain population-wide distributions. This dataset includes samples in five super-population and in 26 populations which a user can view separately or download together. We have made this curated set of disease causing STRs expansions available as a public resource which will be maintained and updated with new disease loci. We believe utilization of the population information can help a user to determine whether the genotyping results could be clinically important.

## Results

To validate and assess STRipy’s performance, we conducted series of simulations where we validated one locus in each gene. First of all, we simulated heterozygous samples by using ART next-generation sequencing read simulator tool [17] whereby the shorter allele was set to be 60 bp in length and varied the length of the longer allele in the range of 60–2100 bp by adding one motif length for each simulation. This includes a large range of repeat lengths and includes the pathogenic cut-off in ~90% of loci. In the same manner, we simulated rarer, expanded homozygous alleles with both alleles the same length ranging from 60 to 2100 bp. These simulations were conducted for each of the 52 genes/loci to examine genotyping accuracy with increasing length of repeats (further details of the methods in S1 Text A; Supplementary information). Once all simulated data sets were generated (63,580 samples), we aligned the simulated reads to the hg38 reference genome using BWA-MEM [18] and genotyped all samples with STRipy using the latest release of the ExpansionHunter (v4.0.2). To estimate the accuracy of calls across simulations we calculated root mean square error (RMSE) for each locus across the different lengths of simulations to determine the genotyping bias (Fig 3).

**Fig 3.**
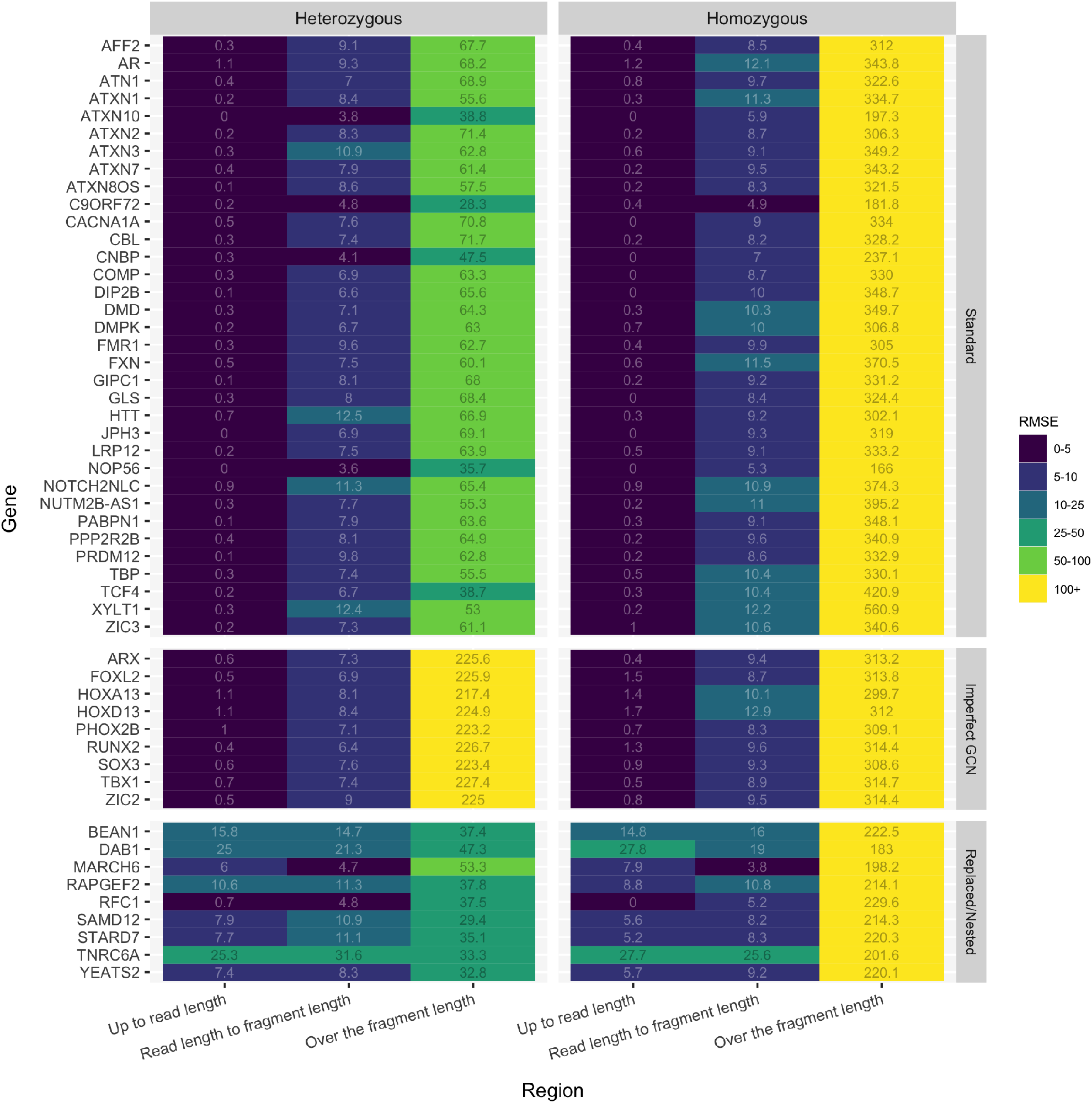
Summary statistics for STRipy’s validation showing root mean square error (RMSE) across simulations in different STR classes and length ranges. “Up to read length” represents samples where the length of repeats is simulated to be between 60 bp to 150 bp, “Read length to fragment length” is repeats from 151 bp to 450 bp and “Over the fragment length” are all simulations where the repeat is between the average fragment length (450 bp) and 2100 bp. RMSE is divided into ranges that is used to colour each cell.

Results were obtained both for homozygous and heterozygous samples at three different STR length ranges considering whether the simulated repeat length is less than the simulated read length of 150 bp, is between the read and the fragment length (mean of 450 bp) or is longer than the fragment length (up to the simulated maximum of 2100 bp). Very low overall RMSE was seen across all standard and imperfect GCN type of repeats ‘up to read length’ in both heterozygous and homozygous samples (median RMSE 0.25 for heterozygous and 0.75 for homozygous samples). We also saw low error rates for genotyping these types of STRs between read and fragment length where the median RMSE for standard repeats is 8.60 and for imperfect GCNs 8.53. Collectively, these results showing very good estimation of alleles up to the fragment length.

We demonstrate the ability to genotype long heterozygous alleles in standard type group that are over the average fragment length (450 bp) and up to 2100 bp, resulting in the median RMSE of 63.17. An example can be seen on Fig 4B showing that STRipy is able to find vast majority of misaligned reads and provide their location to ExpansionHunter as off-target regions, to improve genotyping of long alleles. However, we do see slight underestimation of long alleles which could be due to some fully repeated reads containing more sequencing errors than allowed by STRipy’s algorithm and therefore not extracting them out. Genotyping the same locus using ExpansionHunter with the default variant catalogue shows the inability to genotype long alleles without using misaligned fully repeated reads in off-target regions (Fig 4A). The median RMSE was higher, 224.95 in the imperfect GCN group (an example on Fig 4D). It is important to note that we simulated an extreme case of this type of repeats where the STR locus included four different alanine coding triples in random order which potentially resulted in many unaligned reads of fully repeated GCN sequences and therefore limiting the genotyping of long alleles. We expect more accurate estimation of genotypes in real life samples where the expanded alleles are more uniform and aligned on the genome.

**Fig 4.**
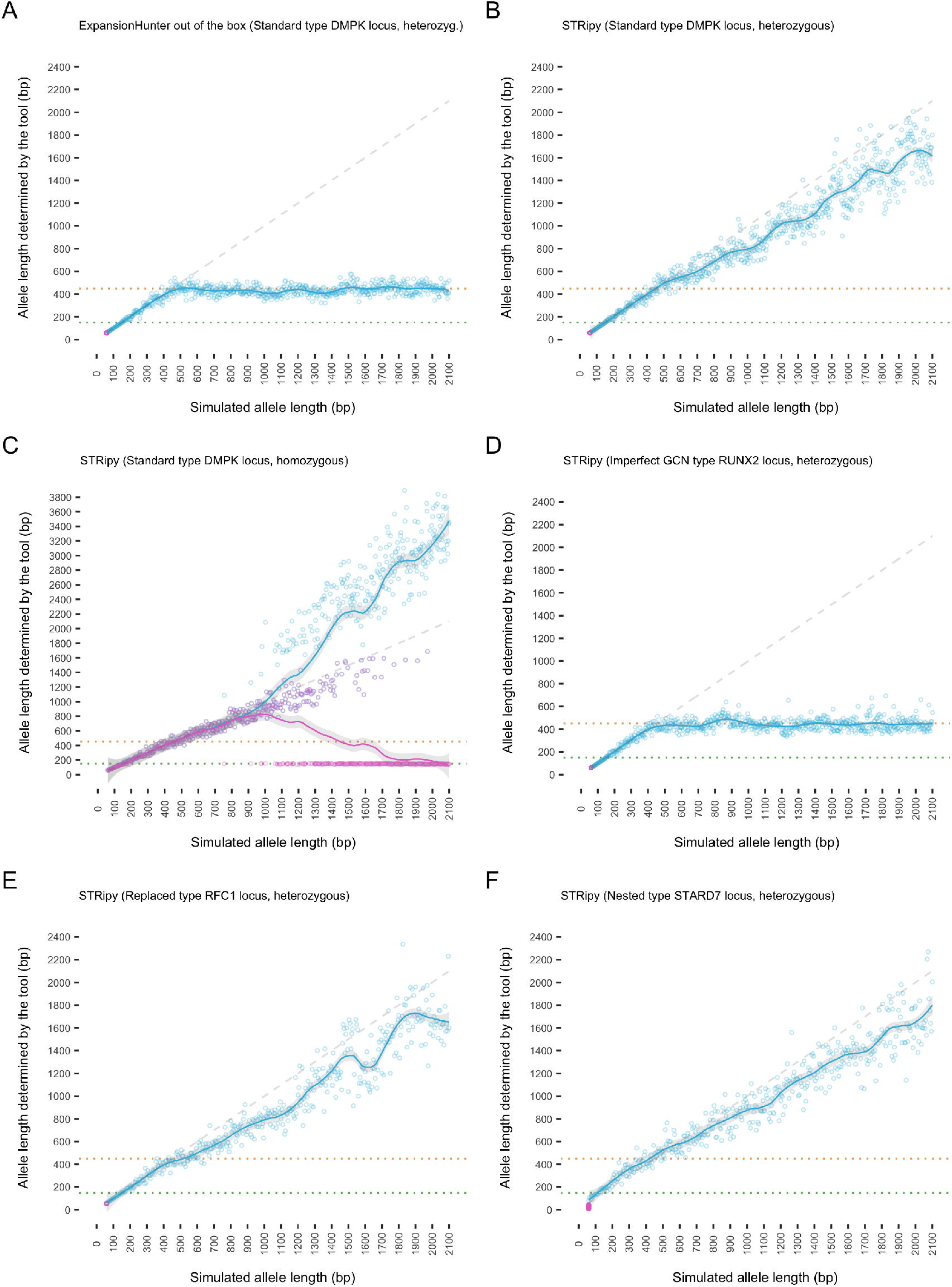
Genotyping results of different types of loci. Pink dots represent one allele, which for heterozygous samples is fixed to 60 bp (bottom left), whereas blue dots represent the other allele that has an increasing number of repeats. Purple dots are overlaps of pink and blue ones. Both alleles were simulated with the same length for homozygous samples. The dotted green line represents the read length and orange one the average fragment length. (A) Results obtained by using ExpansionHunter out of the box with the provided catalogue compared to (B) STRipy’s results which determined off-target regions on the fly allowing genotyping of alleles longer than the fragment length. (C) Example of genotyping issues for long homozygous alleles. (D) Example of genotyping imperfect GCN type repeats, showing slight increase in long alleles compared to A. (E) Example of replaced type repeats (RFC1 gene where biallelic expansions are known to cause cerebellar ataxia, neuropathy, and vestibular areflexia syndrome). (F) Example of nested type repeats (STARD7 locus where expansions are known to cause familial adult myoclonic epilepsy 2).

We saw significantly higher median RMSE (332.06 for standard and 313.20 for imperfect GCN types) when genotyping homozygous samples containing alleles longer than the fragment length. Upon further investigation, we discovered this to be a result of ExpansionHunter algorithm where, from about 700–800 bp, the allele length estimates start to diverge giving heterozygous estimates with one allele an underestimate and one an overestimate (Fig 4C). As we genotyped our samples well beyond that range, the resulting error rate was high. Estimation of alleles over ~700 bp is likely to be error prone with ExpansionHunter v4.0.2 and therefore also in STRipy until this gets resolved in ExpansionHunter.

Regarding the replaced and nested type of repeats, we saw similar results for RFC1 (the only replaced type sample) to the results previously described for standard type repeats (Fig 4E). However, the RMSE in the ‘up to the fragment length’ section was higher for nested type of repeats due to the presence of endogenous repeats which ExpansionHunter cannot distinguish from the pathogenic ones and therefore reported a genotype of both repeat lengths combined (an example in Fig 4F). As we simulated variable numbers of endogenous repeats before and/or after the pathogenic locus, STRipy returned a genotype longer than the simulated insertion which was clearly visible in the ‘up to read length’ range. We developed a feature in STRipy to tackle this issue by counting the number of pathogenic motifs in all spanning/flanking and fully repeated reads. If no pathogenic repeats were found in any of the spanning/flanking or fully repeated reads then this is reported on the results page which indicates the presence of only endogenous repeats. In the vast majority of cases STRipy found fully repeated reads in all samples when the inserted repeated sequence exceeded read length and reported the number of reads to user as an indication of true pathogenic expansions.

Finally, STRipy was applied to a set of whole genome sequenced (WGS) samples that was previously used to validate the genotyping tool STRetch [19] and which contains known pathogenic expansions for nine samples plus one in the intermediate range. We firstly analysed the pathogenic locus in all ten samples (Table 1) in ‘Quick’ mode, for which seven were determined to be in the pathogenic range (five of them with an estimated allele length at least the measured PCR) as well as the one in the intermediate range. For the samples where the allele was less than the PCR length (except number 5), a notification was displayed to the user about a potential presence of a long allele with the recommendation to use ‘Extended’ analysis. When using the ‘Extended’ analysis, the estimated allele length for all affected individuals, except number 5, was in the pathogenic range and for six of them at least the measured PCR value. For the other two, the PCR genotype was in the confidence interval and also close to the estimated one (6 repeat difference). STRipy results and visualisations of sample number 4 are also displayed as an example on the Fig 1.

**Table 1.** STRipy’s performance on true positive whole genome samples.

| # | Gene | Pathogenic cut-off | Measured PCR | STRipy (Quick mode) |  | STRipy (Extended mode) |  |
| --- | --- | --- | --- | --- | --- | --- | --- |
|  |  |  |  | Genotype | CI | Genotype | CI |
| 1 | ATXN1 | 39 | 51 | 30/59 | 30-30/52-69 | 30/59 | 30-30/52-69 |
| 2 | ATXN1 | 39 | 29/32 | 29/34 | 29-29/34-34 | 29/34 | 29-29/34-34 |
| 3 | ATXN3 | 56 | 73 | 24/67 | 24-24/56-82 | 24/67 | 24-24/57-83 |
| 4 | CACNA1A | 21 | 22 | 11/22 | 11-11/22-22 | 11/22 | 11-11/22-22 |
| 5 | AR | 40 | 41 | 36/36 | 27-43/35-52 | 36/36 | 28-44/35-53 |
| 6 | C9orf72 | 31 | >50 | 12/54 | 12-12/41-72 | 12/94 | 12-12/72-127 |
| 7 | CNBP | 55 | >75 | 15/44 | 15-15/41-62 | 15/69 | 15-15/69-127 |
| 8 | DMPK | 50 | >150 | 5/129 | 5-5/98-154 | 5/342 | 5-5/365-407 |
| 9 | AR | 40 | 47 | 50/56 | 38-64/50-81 | 50/56 | 38-64/50-81 |
| 10 | FXN | 70 | ~850 | 115/115 | 62-123/80-156 | 50/871 | 49-89/539-966 |

CI – confidence interval

## Discussion

Here we present STRipy, a software and database that can be easily used to genotype any known disease-causing STRs locus. STRipy works very well to estimate the repeat length in all standard and imperfect GCN type of repeats up to the fragment length as well as standard repeats above the fragment length. This enables accurate genotyping of alleles in the pathogenic range for both standard and imperfect GCN types as the pathogenic range for all GCN type of repeats found so far has not exceeded 40 repeats or 120 bp which is below modern read lengths of 150 bp.

STRipy’s backend uses ExpansionHunter as the genotyping tool and we identified three issues during our study. Firstly, ExpansionHunter exhibits difficulties in correctly genotyping long homozygous alleles above a certain threshold, in our case around 700–800 bp. However, now this has been identified the issue can be mitigated by further manual investigation of the sequencing data to determine the diagnosis of a patient with such long alleles. Out of all the known diseases, this could return an inaccurate genotype (where one allele is not in the pathogenic range) for three diseases – FRAXE mental retardation, glutaminase deficiency with impaired intellectual development and progressive ataxia (GD) and CANVAS, which are all caused by long biallelic expansions exceeding 600 bp. Despite that, a user will be informed of one very long allele which will indicate the need for a follow up analysis. Once this issue is resolved in the ExpansionHunter software, it will be fixed automatically in STRipy as well without the need for changing the code.

Secondly, we observed that the genotyping of nested repeats have higher error rates which is due to ExpansionHunter’s inability differentiate between the pathogenic and endogenous repeat units and therefore reports the sum of these repeats as the length of the genotype. Hence, STRipy is designed to report the number of pathogenic repeat units found in spanning/flanking reads and/or the number of reads fully made of the pathogenic repeat, which should be taken into account when genotyping these loci. The presence of pathogenic repeat units can also be confirmed when examining the read visualisations. Moreover, read visualisations provide information about the sequence in the STR locus, including any interruptions or nucleotide variations that can be present and making an easy visual way for user to understand the STR locus.

Thirdly, during our analysis we often observed that ExpansionHunter detected pathogenic genotypes in the first and second tract of the HOXA13 gene and in some cases, also in tracts of the ARX gene. These are highly conservative sequences and have never been shown to expand as much as we saw. This indicates that these results are false positives and we assume this could be due to the proximity of the two tracts of same, imperfect GCN type motifs. For example, HOXA13_1 and HOXA13_2 both have GCN repeats 63 bp apart and ARX_1 and ARX_2 have same repeats with an 84 bp gap. In STRipy, we are tackling this issue by limiting the region for extracting reads from these loci to 50 bp, which then only extracts out spanning reads in this locus/tract. This results in only using the local reads to infer the genotype with repeats in the adjacent tract not included leading to more accurate results for these loci in STRipy (e.g. HOXA13 loci genotyping shown in 10 samples in S1 Table 5; Supplementary information). Spanning reads are sufficient to genotype all these loci as the highest repeat length discovered so far has been only 30 repeats or 90 bp.

Finally, we improved genotyping of alleles longer than the fragment length by determining off-target regions for misaligned reads from the data itself. However, it is important to note the possibility of observing an overestimated genotype if there are more loci present in the sample, besides the targeted one, that contains long uninterrupted repeats with the same motif and therefore, resulting in fully-repeated reads which are mistakenly used to estimate a genotype.

Here we used simulations to evaluate STRipy, and additionally we genotyped eight different samples with disease causing expansions sequenced with WGS. Diseases caused by STR expansions are rare and some of them have only been found in single families or even individuals which makes it nearly impossible to obtain sequencing data to validate all disease loci using real biological samples. Realistically, we can validate all loci only by using simulations and this is the main limitation of our validation. Data from real sequenced samples might be noisy and might be more complex to analyse than simulated samples. However, we created a robust and comprehensive simulation dataset to more closely resemble real life data by introducing variability in the fragment size, simulated reads with natural occurring sequencing errors and created alleles for replaced/nested type of repeats that match with biological findings.

STRipy fills a specific niche in the field and is meant to serve as the first line screening tool for STR diseases that is available to the wider community. STRipy allows the targeting of one locus in one sample at the moment and therefore, it is more a complementary tool for ExpansionHunter and other similar tools developed that can be incorporated into bioinformatics pipelines and used in large scale multi-loci analysis. We have also provided a tool for creating ExpansionHunter’s variant catalogue to assist users with such large-scale analyses, which contains all loci described in this paper.

STRipy and the STRs database is available at https://STRipy.org. As new disease causing STR expansions are discovered these will be added to STRipy’s database and genotyping functionality.

## Data availability

Source code for STRipy’s Client can be retrieved at https://gitlab.com/andreassh/stripy-client and for the STRipy’s Server at https://gitlab.com/andreassh/stripy-server. All other code and data used in this paper is available on Zenodo (doi: 10.5281/zenodo.4939980). The true positive biological whole genome samples are available from the Sequence Read Archive under the accession number SRP148723 (individual samples accessions are from SRX4114164 to SRX4114173).

