## Supplementary information for "STRipy: a graphical application for enhanced genotyping of pathogenic short tandem repeats in sequencing data"

S1 Table 1. All known disease-causing STR loci made of standard type of repeats.

| Gene | Reference and location | Motif | Disease | Inheritance | Normal range | Intermediate range | Pathogenic cut-off | References |
|---|---|---|---|---|---|---|---|---|
| AFF2 (FMR2) | chrX:148500638-148500683 (5′ UTR) | CCG | FRAXE mental retardation (FRAXE) | XLR | 6–25 | – | ≥200 | [1] |
| AR | chrX:67545316-67545385 (Coding) | CAG | Spinal and bulbar muscular atrophy (SBMA) | XLR | 17–35 | – | ≥40 | [2–4] |
| ATN1 | chr12:6936728-6936773 (Coding) | CAG | Dentatorubral-pallidoluysian atrophy (DRPLA) | AD | 7–23 | – | ≥49 | [5,6] |
| ATXN1 | chr6:16327635-16327722 (Coding) | CAG | Spinocerebellar ataxia 1 (SCA1) | AD | 6–32 | 36–38 | ≥39 | [7,8] |
| ATXN2 | chr12:111598950-111599019 (Coding) | CAG | Spinocerebellar ataxia 2 (SCA2) | AD | 13–31 | 32–34 | ≥35 | [9–11] |
| ATXN3 | chr14:92071010-92071040 (Coding) | CAG | Spinocerebellar ataxia 3 (SCA3) | AD | 12–44 | 45–55 | ≥56 | [12,13] |
| ATXN7 | chr3:63912685-63912715 (Coding) | CAG | Spinocerebellar ataxia 7 (SCA7) | AD | 4–35 | 28–35 | ≥34 or ≥36 | [14,15] |
| ATXN8OS | chr13:70139383-70139428 (3′ UTR) | CTG | Spinocerebellar ataxia 8 (SCA8) | AD | 2–37 | 38–79$ | ≥80 | [16–18] |
| ATXN10 | chr22:45795354-45795424 (Intron) | ATTCT | Spinocerebellar ataxia 10 (SCA10) | AD | 10–29 | 30–799$ | ≥800 | [19] |
| C9orf72 | chr9:27573528-27573546 (Intron) | GGGGCC | Amyotrophic lateral sclerosis and/or frontotemporal dementia (FTDALS1) | AD | 1–23 | 20–30* | ≥31* | [20,21] |
| CACNA1A | chr19:13207858-13207897 (Coding) | CAG | Spinocerebellar ataxia 6 (SCA6) | AD | 4–18 | 19 | ≥21 | [22–24] |
| CBL | chr11:119206289-119206322 (5′ UTR) | CCG | Jacobsen syndrome (JBS) | NI | 8–80 | 85–100 | ≥101 | [25] |
| CNBP (ZNF9) | chr3:129172576-129172656 (Intron) | CCTG | Myotonic dystrophy 2 (DM2) | AD | ≤29 | 30–54 | ≥55 | [26,27] |
| COMP | chr19:18786034-18786049 (Coding) | GAC | Multiple epiphyseal dysplasia (MED) | AD | 5 | – | 6 | [28] |
| | | | Pseudoachondroplasia (PSACH) | AD | 5 | – | 4 & 7 | [28,29] |
| DIP2B | chr12:50505003-50505024 (5′ UTR) | CGG | FRA12A mental retardation (MRFRA12A) | AD | 6–23 | – | ≥270 | [30] |

| | | | | | | | | |
|---|---|---|---|---|---|---|---|---|
| DMD | chrX:31284557-31284605 (Intron) | GAA | Duchenne muscular dystrophy (DMD) | XLR | 11–33 | – | ≥59 | [31] |
| DMPK | chr19:45770204-45770264 (3′ UTR) | CTG | Myotonic dystrophy 1 (DM1) | AD | 5–34 | 35–49 | ≥50 | [32–34] |
| FMR1 | chrX:147912050-147912110 (5′ UTR) | CGG | Fragile X Syndrome (FXS) | XLD | 6–40 | 41–54 | ≥201 | [35,36] |
| | | | Fragile X tremor/ataxia syndrome (FXTAS) | XLD | 6–40 | 41–54 | ≥55 ≤200 | [35,36] |
| FXN | chr9:69037286-69037304 (Intron) | GAA | Friedreich ataxia (FRDA) | AR | 5–30 | – | ≥70 | [37] |
| GIPC1 | chr19:14496041-14496074 (5′ UTR) | GGC | Oculopharyngodistal myopathy 1 (OPDM2) | AD | 12–32 | – | ≥73 | [38] |
| GLS | chr2:190880872-190880920 (5′ UTR) | GCA | Glutaminase deficiency GD) | AR | 5–26 | – | ≥680 | [39] |
| HTT | chr4:3074876-3074933 (Coding) | CAG | Huntington's disease (HD) | AD | 9–26 | 27–35 | ≥36 | [40–42] |
| JPH3 | chr16:87604287-87604329 (3′ UTR) | CTG | Huntington disease-like 2 (HDL2) | AD | 6–27 | – | ≥41 | [43] |
| LRP12 | chr1:149390805-149390831 (5′ UTR) | CGG | Oculopharyngodistal myopathy 1 (OPDM1) | AD | 13–45 | – | ≥93 | [44] |
| NOP56 | chr20:2652733-2652757 (Intron) | GGCCTG | Spinocerebellar ataxia 36 (SCA36) | AD | 3–14 | – | ≥650 | [45,46] |
| NOTCH2NLC (NBPF19) | chr1:149390802-149390829 (5′ UTR) | GGC | Neuronal intranuclear inclusion disease (NIID) | AD | 7–39 | – | ≥90 | [44] |
| | | | Hereditary essential tremor type 6 (ETM6) | AD | 4–41 | – | ≥60 | [47] |
| NUTM2B-AS1 | chr10:79826377-79826403 (Intron) | CGG | Oculopharyngeal myopathy with leukoencephalopathy 1 (OPML1) | AD | 3–16 | – | ≥700 | [44] |
| PABPN1 | chr14:23321472-23321490 (Coding) | GCG | Oculopharyngeal muscular dystrophy (OPMD) | AD or AR | 6 | – | ≥7 | [48] |
| PPP2R2B | chr5:146878728-146878758 (5′ UTR) | CAG | Spinocerebellar ataxia 12 (SCA12) | AD | 7–31 | – | ≥51 | [49,50] |
| PRDM12 | chr9:130681606-130681639 (Coding) | CCG | Neuropathy, hereditary sensory and autonomic, type VIII (HSAN8) | AR | ≤14 | – | ≥18 | [51] |
| TBP | chr6:170561907-170562015 (Coding) | CAG | Spinocerebellar ataxia 17 (SCA17) | AD | 25–42 | – | ≥43 | [52,53] |

| | | | | | | | | |
|---|---|---|---|---|---|---|---|---|
| TCF4 | chr18:55586155-55586227 (Intron) | CTG | Fuchs endothelial corneal dystrophy-3 (FECD3) | AD | ≤31 | – | ≥50 | [54,55] |
| XYLT1 | chr16:17470907-17470922 (5′ UTR) | GGC | Desbuquois dysplasia-2 (DBQD2) | AR | 9–20 | – | ≥110 | [56] |
| ZIC3 | chrX:137566826-137566856 (Coding) | GCC | X-linked VACTERL syndrome (VACTERLX) | XLR | 10 | – | ≥12 | [57] |

The coordinates are in hg38 assembly and 0-based. Asterisk (*) denotes data where the range vary in literature. $Denotes a range specified by us based on the findings in literature from multiple sources. Normal, intermediate and pathogenic ranges were derived from the articles marked in the references field. AD – autosomal dominant, AR – autosomal recessive, XLD – X-linked dominant, XLR – X-linked recessive, NI – not inherited.

S1 Table 2. All known disease-causing STR loci made of imperfect GCN type of repeats.

| **Gene** | **Reference and location** | **Motif** | **Disease** | **Inheritance** | **Normal range** | **Intermediate range** | **Pathogenic cut-off** | **References** |
|---|---|---|---|---|---|---|---|---|
| ARX | †chrX:25013529-25013565<br>‡chrX:25013649-25013697<br>(Coding) | GCN | Developmental and epileptic encephalopathy-1 (DEE1) | XLR | 12†<br>16‡ | – | ≥20†<br>≥23‡ | [58] |
| | | | Partington syndrome (PRTS) | XLR | 12† | – | ≥20† | [58] |
| | | | X-linked mental retardation with or without seizures (MRXARX) | XLR | 12†<br>16‡ | – | ≥20†<br>≥18‡ | [59] |
| FOXL2 | chr3:138946020-138946062 (Coding) | GCN | Blepharophimosis, ptosis, and epicanthus inversus syndrome (BPES) | AD | 14 | – | ≥19 | [60,61] |
| HOXA13 | +chr7:27199924-27199966<br>†chr7:27199825-27199861<br>‡chr7:27199678-27199732<br>(Coding) | GCN | Hand-foot-genital syndrome (HFG) | AD | 14+<br>12†<br>18‡ | – | ≥22+<br>≥18†<br>≥24‡ | [62–64] |
| HOXD13 | chr2:176093058-176093103 (Coding) | GCN | Synpolydactyly (SPD) | AD | 15 | – | ≥22 | [65] |
| PHOX2B | chr4:41745971-41746031 (Coding) | GCN | Central hypoventilation syndrome (CCHS) | AD | 20 | – | ≥24 | [66] |
| RUNX2 | chr6:45422750-45422801 (Coding) | GCN | Cleidocranial dysplasia (CCD) | AD | 17 | – | ≥27 | [67] |
| SOX3 | chrX:140504316-140504361 (Coding) | GCN | X-linked mental retardation (XLMR) | XLR | 15 | – | ≥26 | [68] |
| | | | X-linked panhypopituitarism (PHPX) | XLR | 15 | – | ≥22 | [69] |
| TBX1 | chr22:19766762-19766807 (Coding) | GCN | Tetralogy of Fallot (TOF) | AD | 15 | – | ≥25 | [70] |
| ZIC2 | chr13:99985448-99985493 (Coding) | GCN | Holoprosencephaly 5 (HPE5) | AD | 15 | – | ≥25 | [71] |

Given coordinates are in hg38 assembly and 0-based. +, † and ‡ are used to distinguish values for different tracts in a locus. Normal, intermediate and pathogenic ranges were derived from the articles marked in the references field. AD – autosomal dominant, XLR – X-linked recessive.

S1 Table 3. All known disease-causing STR loci made of nested and replaced types of repeats.

| Gene | Reference and location | Motif | Disease | Inheritance | Normal range | Intermediate range | Pathogenic cut-off | References |
|---|---|---|---|---|---|---|---|---|
| BEAN1 | chr16:66490398-66490466 (Intron) | TGGAA *(TAAAA)* | Spinocerebellar ataxia 31 (SCA31) | AD | 0 | – | ≥500# | [72] |
| DAB1 | chr1:57367043-57367118 (Intron) | ATTTC *(ATTTT)* | Spinocerebellar ataxia 37 (SCA37) | AD | 0 | – | ≥31 | [73,74] |
| RFC1 | chr4:39348424-39348485 (Intron) | AAGGG *(AAAAG)* | Cerebellar ataxia, neuropathy, and vestibular areflexia syndrome (CANVAS) | AR | 0§ | – | ≥400 | [75,76] |
| SAMD12 | chr8:118366816-118366914 (Intron) | TTTCA *(TTTTA)* | Familial adult myoclonic epilepsy 1 (FAME1) | AD | 0 | – | ≥100 | [77–79] |
| STARD7 | chr2:96197066-96197121 (Intron) | TTTCA *(TTTTA)* | Familial adult myoclonic epilepsy 2 (FAME2) | AD | 0 | – | ≥274 | [80] |
| MARCH6 | chr5:10356347-10356407 (Intron) | TTTCA *(TTTTA)* | Familial adult myoclonic epilepsy 3 (FAME3) | AD | 0 | – | ≥668# | [81] |
| YEATS2 | chr3:183712187-183712222 (Intron) | TTTCA *(TTTTA)* | Familial adult myoclonic epilepsy 4 (FAME4) | AD | 0 | – | ≥1000# | [82] |
| TNRC6A | chr16:24613439-24613529 (Intron) | TTTCA *(TTTTA)* | Familial adult myoclonic epilepsy 6 (FAME6) | AD | 0 | – | ≥1100# | [78] |
| RAPGEF2 | chr4:159342526-159342616 (Intron) | TTTCA *(TTTTA)* | Familial adult myoclonic epilepsy 7 (FAME7) | AD | 0 | – | ≥60 | [78,79] |

The coordinates are in hg38 assembly and 0-based. Coordinates for the reference region denotes the STR location of endogenous repeats. The motif field indicates the pathogenic motif with the non-pathogenic reference motif in parentheses. Normal, intermediate and pathogenic ranges were derived from the articles marked in the references field. §The normal range is set to 0 because the pathogenic motif has not been found in the healthy population, except in one alleles of the CANVAS patients. #Size of the whole allele (including non-pathogenic repeats). Number of repeats in the pathogenic cut-off column for many diseases has been calculated from the reported allele size assuming they are composed solely of pentanucleotide repeats. AD – autosomal dominant, AR – autosomal recessive.

**Text 1. Methods used for validation of STRipy and population-wide genotyping.** This section describes methods used for validating STRipy and creating the STRs database.

**Method of simulating samples for validation.**

Creation of samples with artificial length of repeats was done in two main steps. Firstly, a template sequence was created based on the reference genome, where the known STR sequence was swapped with a newly created one. The start and end positions of the STR sequence were derived from the literature, assessed manually and in some cases the end position was adjusted in a way that it ends before at least two consecutive repeats of the non-pathogenic motif was present (coordinates shown in S1 Table 1–3). Secondly, this template was used to simulate reads that were then aligned to the reference genome.

We wrote a custom-made program that takes the genomic coordinates of a known pathogenic STR locus and extracts out 2 kb flanking sequence before and after the repeated sequence. Then, a new stretch of repeats was created with specific length, which was then merged with the flanking sequences to form over a 4 kb sequence and saved into a new FASTA file. For all the disorders that are caused by imperfect GCN types of repeats (stretches of the same amino acid but with an inconsistent repeat sequence coding alanine), we generated stretches of motifs that had one base randomly chosen. For example, to simulate polyalanine stretches we used GCN as the motif and replaced "N" randomly with either "A", "T", "G" or "C" in every repeat present in the sequence.

To determine genotyping accuracy across increasing length of repeats, we first simulated homozygous alleles that included the pathogenic motif from 60 bp until the 2100 bp limit was reached where each simulation increased the length of the allele by one repeat unit (in the range of 20–700 trinucleotide-, 15–525 tetranucleotide-, 12–420 pentanucleotide and 10–350 hexanucleotide repeats). In the case of the nested type of repeats, we created short alleles without the pathogenic motif. Secondly, for heterozygous alleles we designed the shorter allele to be fixed length (60 bp) and increased the length of the second allele from 60 bp until 2100 bp, similarly to the process of simulating homozygous repeats. For alleles of the nested type of repeats, we simulated alleles based on the formulas in S1 Table 4, where *exp* for the non-pathogenic repeat unit was a random number between 2 and 10 and for pathogenic motif the repeat was in increasing length. For example, to simulate FAME4 disorder, a normal AAAAT repeats was created which length was randomly chosen to be between 2 and 10 repeats, followed by 12 to 240 repeats of pathogenic AAATG motif to create a 60 to 2100 bp pathogenic locus.

S1 Table 4. Formulas used to simulate alleles of replaced and nested type repeats.

| **Gene** | **Disease** | **Repeat type** | **Formula for allele simulation** |
|---|---|---|---|
| RFC1 | CANVAS | Replaced | $(AAAAG)_{11}[AAGGG]_{exp}$ |
| BEAN1 | SCA31 | Nested | $(TCAC)_{1}[TGGAA]_{exp}(TAGAA)_{exp}(TAAAATAGAA)_{exp}$ |
| DAB1 | SCA37 | Nested | $(AAAAT)_{60\text{-}79}[GAAAT]_{31\text{-}75}(AAAAT)_{58\text{-}90}$ |
| MARCH6 | FAME1 | Nested | $(TTTTA)_{12}[TTTCA]_{exp}$ |
| RAPGEF2 | FAME2 | Nested | $(TTTTA)_{exp}[TTTCA]_{exp}(TTTTA)_{exp}$ |
| SAMD12 | FAME3 | Nested | $(AAAAT)_{exp}[TGAAA]_{exp}$ |
| STARD7 | FAME4 | Nested | $(AAAAT)_{exp}[AAATG]_{exp}$ |
| TNRC6A | FAME6 | Nested | $(TTTTA)_{22}[TTTCA]_{exp}(TTTTA)_{exp}$ |
| YEATS2 | FAME7 | Nested | $(TTTTA)_{exp}[TTTCA]_{exp}$ |

Motif in parentheses is the non-disease causing and motif in square brackets is the pathogenic one. The number after parentheses or square brackets shows the number of repeats of that motif, exp denotes an expansion which length is not specified and will be randomly assigned.

We generated paired-end reads in the FASTQ format from the template (FASTA) files with the ART (version MountRainier 2016-06-05) next-generation sequencing read simulator tool [83]. Illumina HiSeqX PCR-free profile was used to simulate 150 bp reads with the mean fragment size of 450 bp and standard deviation 50 bp. Sequences were simulated to have total coverage of 50x, both for homozygous and heterozygous samples. All FASTQ files were aligned on the UCSC GRCh38.p12 reference genome with BWA-MEM v0.7.17 [84] and indexed with Samtools v1.10 [85].

### Genotyping samples with STRipy

We used STRipy with ExpansionHunter v4.0.2 as the genotyper. STRipy was performed on each of the aligned sample (both homozygous and heterozygous) and results saved into a separate file that was analysed with R (paper_analysis.R) [86]. We ran STRipy by using the default parameters, except switching on the option to use alternative contigs.

We used root mean square error (RMSE) to determine genotyping accuracy for both, heterozygous and homozygous samples by comparing predicted and true repeat lengths as described by Mousavi et al. (2019). The $\langle x_1^i, x_2^i \rangle$ denotes a diploid genotype of sample $i$ and is ordered by length such that $x_1^i \leq x_2^i$. To compare the actual $X = \{\langle x_1^1, x_2^1 \rangle, \langle x_1^2, x_2^2 \rangle \ldots \langle x_1^n, x_2^n \rangle\}$

and predicted $Y = \{\langle y_1^1, y_2^1\rangle, \langle y_1^2, y_2^2\rangle \ldots \langle y_1^n, y_2^n\rangle\}$ genotypes, then RMSE is defined by the following formula [87]:

$$\sqrt{\sum_{i=1}^{n} \sum_{j=1}^{2} \frac{(y_j^i - x_j^i)^2}{2n}}$$

**Population-wide data creation**

We used 2,504 samples from the DRAGEN reanalysis of the 1000 Genomes Dataset (1kGP-DRAGEN, https://registry.opendata.aws/ilmn-dragen-1kgp) to genotype all known pathogenic loci in all samples. These samples were sequenced with the Illumina NovaSeq 6000 system to produce 150 bp paired-end reads of at least 30x coverage. All BAM files in the dataset were realigned to hg38 assembly by using the Illumina DRAGEN v3.5.7b (https://www.illumina.com/products/by-type/informatics-products/dragen-bio-it-platform.html).

All samples in Amazon Web Services S3 instance: s3://1000genomes-dragen/data/dragen-3.5.7b/hg38_altaware_nohla-cnv-anchored were analysed. ExpansionHunter v4.0.2, with our complete variant catalogue that included all genomic loci, including all tracts in genes. We applied a filter to exclude results where there are less than 5 spanning reads. In case of no spanning reads, then the criterion was minimum of 50 combined flanking and in-repeat reads to increase confidence of determined genotypes. To genotype replaced and nested types of repeat loci, we created a custom-made script (genotype-nested.py) [86] which analysed spanning reads in the realigned BAM files outputted from ExpansionHunter and determined the length of pathogenic repeats in the these reads by using previously described genotyping model [88].

Next, we determined all samples where the allele was over the read length of these samples and genotyped these samples again by using a catalogue that included pre-defined off-target regions to enable genotyping long alleles. To do this, we simulated one sample for each locus whose coverage was increased to 100x and we shifted down the quality score of every read by 10 to increase sequencing error rate to 1% of the default profile. Following alignment, we determined all regions where reads aligned to and merged overlapping and nearby regions within 100 bp into one single entry and adding 50 bp on both ends. These coordinates were specified as off-target regions in the variant catalogue. Off-target regions that included or overlapped with the reference region of the disease or regions on alternative contigs were removed from the list beforehand.

S1 Table 5. Genotypes of three tracts in HOXA13 gene in the ten real biological samples estimated by ExpansionHunter and STRipy.

| | ExpansionHunter | | | STRipy | | |
|---|---|---|---|---|---|---|
| # | HOXA13_1 | HOXA13_2 | HOXA13_3 | HOXA13_1 | HOXA13_2 | HOXA13_3 |
| 1 | 14/14 | 12/12 | 18/18 | 14/14 | 12/12 | 18/18 |
| 2 | 14/14 | 12/12 | 18/18 | 14/14 | 12/12 | 18/18 |
| 3 | 14/14 | 12/66 | 18/18 | 14/14 | 12/12 | 18/18 |
| 4 | 14/66 | 12/70 | 18/18 | 14/14 | 12/12 | 18/18 |
| 5 | 14/14 | 12/12 | 18/18 | 14/14 | 12/12 | 18/18 |
| 6 | 14/14 | 12/12 | 18/18 | 14/14 | 12/12 | 18/18 |
| 7 | 14/80 | 12/73 | 18/18 | 14/14 | 12/12 | 18/18 |
| 8 | 14/14 | 12/12 | 18/18 | 14/14 | 12/12 | 18/18 |
| 9 | 14/69 | 12/12 | 18/18 | 14/14 | 12/12 | 18/18 |
| 10 | 14/14 | 12/12 | 18/18 | 14/14 | 12/12 | 18/18 |

### List of references

10.1016/j.nbd.2019.104635

62. Innis JW, Mortlock D, Chen Z, Ludwig M, Williams ME, Williams TM, et al.

Polyalanine expansion in HOXA13: Three new affected families and the molecular consequences in a mouse model. Hum Mol Genet. 2004;13(22):2841–51. doi: 10.1093/hmg/ddh306
